# StomaQuant: Deep Learning-Based Quantification for Stomatal Trait Assessment

**DOI:** 10.64898/2026.01.06.698068

**Authors:** Kenny J.X. Lau, Cheng-Yen Chen, Sahanna Muruganantham, Vayutha Muralishankar, Naweed I. Naqvi

**Author notes:** Corresponding Author: Naweed I. Naqvi or Kenny Lau.

## Abstract

Stomata are microscopic pores that play a vital role in transpiration and gaseous exchange from leaf surfaces in plants. The stomatal density and size directly influence photosynthesis and hydrodynamics capacity. Conventional approaches for counting and determining stomatal density is labour-intensive and lack scalability. Although there are several AI-based stomata finder tools that were published in the last decade, existing models were trained on model plants like wheat, barley and *Arabidopsis*. Stomata in such model plants are generally elliptical, but applying a universal model to all plant species is not feasible due to their diverse morphological characteristics. Previous studies have suggested using the stomatal index to quantify the ratio between epidermal cells and total stomatal count. However, this approach can be difficult to apply consistently, as epidermal cell shape and size vary across plant species. Instead, we propose measuring stomatal density based on the number of stomata per total imaged pixel area in the captured images. In this study, a comparison between YOLOv12 and RF-DETR models were made for real-time stomata detection in normal and difficult-to-image and out-of-focus occluded images. The in-house training dataset consisted of images of 300 rice,100 barley and 50 sugarcane leaves that were captured against a dark background. YOLOv12 outperformed RF-DETR with higher mAP50:95 score. The models were trained with image augmentation for 300 epochs and YOLOv12 achieved a peak mean average precision of 98.5% and exceled at detecting stomata across abaxial and adaxial surfaces of leaves of both monocot and dicot plants. StomaQuant has also been shown to be effective for both epidermal peel and ethanol decolorised samples. Thus, StomaQuant can be used to effectively and efficiently estimate the stomatal density and size in a wide range of host plant species.

## INTRODUCTION

The global population is projected to surpass 9 billion by 2050, concerns about the future of global food security are mounting as climate patterns becoming more unpredictable [1]. Rainfed agricultural cultivation are highly susceptible to variability driven by climate change and water management [2]. Water is lost in plants through transpiration via the stomata. To improve drought tolerance, genetic manipulation can select for mutants with lower stomatal density, and this approach has shown success in improving water-use efficiency in *Arabidopsis thaliana* [3, 4].

Stomata play a vital role in gaseous exchange, especially for carbon dioxide, wherein their density, size and aperture directly influence photosynthesis and transpiration [5]. Stomatal imaging relies heavily on the choice of sample preparation technique, which falls into two categories, namely destructive and non-destructive methods. Non-destructive approaches, such as nail polish imprints or portable microscopy aims to preserve the integrity of the leaf to allow dynamic observation of stomatal behaviour [6]. However, these methods are difficult to reproduce with precision as nail polish imprints can introduce artifacts such as air bubbles or leave partial imprint that obscure fine structures, especially in plants with smaller leaf surface area and smaller stomata size. On the other hand, destructive methods such as ethanol fixation and decolouration offer the advantage of long-term sample preservation and facilitate high-resolution imaging without compromising structural details. However, destructive methods cause damage to the plants and affect *in-vivo* observations of the opening and closing of the stoma making longitudinal monitoring of the plant health unfeasible [7]. Depending on the sample preparation techniques used, downstream artificial intelligence (AI) and computer vision model training can be compromised if image quality is poor due to artifacts and low resolution that may reduce the overall model accuracy.

The stomata for model plants are typically elliptical. In dicots, each stoma is flanked by two kidney-shaped guard cells. In contrast, monocots possess two additional subsidiary cells encircling the guard cells, resulting in a distinctive dumbbell-shaped stoma. The geometry of stomata varies among monocots as the size of subsidiary cells may also differ in different plants. For instance, a stoma in rice has two triangular shaped subsidiary cells that give the stoma an overall rhombus shaped appearance, and this stomatal structure may not be detected in AI models that were specifically trained on elliptical-shaped stomata dataset. Additionally, some studies proposed the use of stomatal index to quantify the ratio between stomata and total stomata with epidermal cell count [8,9]. Stomatal index may be difficult to measure as the shape and size of epidermal cells also vary in different plants.

Previous studies have shown that it is possible to automate the detection and counting of stomata in *Arabidopsis* (dicot) and barley (monocot) [10]. Stomata segmentation is useful in plant biology as it allows for the detailed analysis of stomatal traits that may influence key physiological states through gas exchange, transpiration and photosynthesis. Stomatal analysis is essential for understanding plant responses to environmental stress like drought and optimising yield. In the last decade, the emergence of computer vision and deep learning techniques have transformed stomata segmentation in plant leaf imaging, enabling automated and highly efficient stomata analysis. In particular, the advent of convolutional neural networks (CNNs) marked a transformative shift in the field [11]. One of the pioneer models, Fully Convolutional Networks (FCNs) introduced pixel-level classification by replacing traditional fully connected layers with convolutional ones for end-to-end training on segmentation tasks [12]. Building on the success of FCNs, U-Net was created, and it featured a more advanced encoder-decoder architecture that significantly improved segmentation accuracy in biomedical imaging by integrating fine-grained spatial details [13]. Semantic segmentation models classify every pixel in an image to an object class and excels at object detection whereas instance segmentation models classify each pixel and distinguish the different instances of the same object class facilitating object counting. Instance segmentation models such as mask R-CNN and YOLOv8 have demonstrated exceptional performance across a wide range of object detection applications including stomata counting task with reported mean average precision scores of 0.97920 and 0.98164 respectively [14]. In a separate study involving wheat and poplar, semantic segmentation was employed to predict stomatal density and size to calculate and estimate water loss based on the concept of maximum stomatal conductivity [15]. Interestingly, the MobileNet, based on the CNN algorithm, has also been trained to detect and assess stomatal density for drought-tolerant oil palm breeding [16]. The mask R-CNN model was also used in the StomaAI tool for stomatal localization and phenotypic data collection [10]. RotatedStomataNet uses R^3^ det network to train images from destructive and non-destructive dataset as well as on monocot and dicot plants providing comprehensive analysis [17]. There is a gradual shift from CNN-based models towards transformer-based models, and this is possible with more efficient attention-based transformer models [18]. Transformer-based object detector like the RF-DETR has been demonstrated to outperform CNN-models in instance segmentation tasks achieving higher mAP scores and converges faster with lesser training epochs even on lower specification hardware machines in detecting occluded green fruit [19]. In addition, Meta AI has also launched its cutting edge self-supervised model, Distillation of knowledge with NO labels (DINOv3), that was trained on 1.7 billion images with 7 billion parameters without human annotations, providing it as a foundational model for image segmentation and object detection applications [20].

Obtaining labelled stomatal images from diverse plant species in the world is likely unfeasible due to the vast biological variability and annotation challenges. Transformer-based models excel in learning rich representations from unlabelled images and generalize object of interest effectively. The recent release of RF-DETR by RoboFlow, showed outstanding performance by surpassing mAP score of 60% on the MS COCO dataset and has topped the charts of transformer-based model to-date. In this study, we used both YOLOv12 and RF-DETR models for the task of stomatal detection and density measurement. We also showed that both object detection models can be used for drought tolerant plant breeding program and precision agriculture in barley, rice and sugarcane.

## MATERIALS AND METHODS

### Plant Cultivation and Sample Preparation

Rice (*Oryza sativa*) was grown for 2 weeks after transplantation to five leaf stage. Barley (*Hordeum vulgare*) seedlings were grown for a month. Sugarcane (*Saccharum officinarum*) saplings were grown for 3 months in a greenhouse facility at Lim Chu Kang, Singapore (103⁰70’49’’ E and 1⁰42’73’’ N). The leaves were excised with scissors and immersed in 70% ethanol. The ethanol-soaked leaves were then incubated in a 55°C water bath overnight to decolourize and remove the chlorophyll pigments from the leaves. The decolourized leaves were cut into small squares and placed on a glass slide and cover slip for microscopic imaging. All such rice, barley and sugarcane leaf explants were then imaged using the Olympus BX53 microscope (Evident Scientific, Japan) under the 20× and 40× magnifications at 1392 × 1040 pixels resolution. Data acquisition was performed in Temasek Life Sciences Laboratory, Singapore. A collection of 450 images of abaxial and adaxial surfaces of barley, rice and sugarcane leaves were captured to train the YOLOv12 and RF-DETR models for stomatal detection.

### YOLOv12 Model Training Parameters

The YOLOv12 model was trained using a two-step approach, binary mask segmentation and convolutional neural network. Segmentations were done manually in labelme by labelling the object as stomata, where binary mask in white (RGB: 255, 255, 255) denotes the stomata and black (RGB: 0, 0, 0) the background. A python script was written to convert the binary mask into polygons. Localisation of the polygons were transcribed into JSON format and (You Only Look Once) YOLO txt format. Subsequently, training was then performed on the pair of original images and corresponding annotation files using the YOLOv12s model. Images were randomly split into 80% training and 20% testing. Image augmentation was performed by scaling, flipping vertically or horizontally and rotating (Fig S1). The training parameters were set to AdamW optimizer learning rate of 0.002, momentum of 0.9 and max stride of 1216. Model training was performed on a high-performance computing cluster that has an AMD EPYC 7543P 32-core processor CPU for 300 epochs and batch size of 4. For downstream statistical analysis, unseen images that were not used for training were used for testing. Statistical analyses were performed in R using *ggplot2* [21].

### RF-DETR Model Training Parameters

YOLO annotations were converted into COCO format. The RF-DETR (Roboflow Detection Transformer) was trained using the same set of images that were used with YOLO for training, validation and testing with 80%-15%-5% ratio. Unlike YOLOv12, the RF-DETR tool does not have a built-in augmentation pipeline, hence augmentation was performed externally using a python script. The script is available at Dryad repository <u>10.5061/dryad.3j9kd51zr</u>. Augmentation parameters include scaling, flipping and rotation. Model training was performed on a high-performance computing cluster that has an AMD EPYC 7543P 32-core processor CPU for 300 epochs and batch size of 4.

### Evaluation Metrics

**True Positive (TP)**: A predicted bounding box correctly identifies a ground truth stoma with an Intersection over Union (IOU) greater than 0.5 threshold.

**False Positive (FP)**: A predicted bounding box that incorrectly overlap with the ground truth box with IOU less than 0.5.

**False Negative (FN)**: Stoma that is overlooked by the detection model with no predicted bounding box.

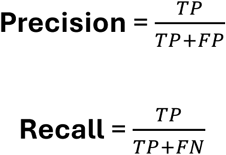

Intersection over Union (IoU)

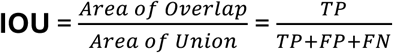

**mAP50** measures the mean Average Precision at IOU threshold of 0.50 whereas

**mAP50-95** splits the Average Precision into ten IOU levels from 0.50 to 0.95 with increments of 0.05 to detect if predicted bounding box strictly overlaps the stoma of interest.

**Precision** measures the proportion of predicted bounding boxes that are correct

**Recall** measures the proportion of actual objects that model successfully detects

## RESULTS

The results for stomata detection in barley, rice and sugarcane leaves with YOLOv12 are as shown in Figure 1. Top panel shows the evaluation metrics for training images while the bottom panel for validation images. Box loss refers to getting the location of the bounding box in the correct position with the least error. Class loss refers to classifying stomata and the background correctly. Distribution Focal Loss (DFL) is a loss function for improving object detection accuracy by reducing the weight of easy-to-detect instances and focuses on hard-to-detect objects. The model converged after epoch 28 at mAP50 of 0.90 and eventually reached mAP50 of 0.95 at epoch 45. After 300 epochs of training, the final model achieved a peak mAP50 of 0.98, mAP50-95 of 0.77 and recall of 0.97. The accuracy of the YOLOv12 model was evaluated based on the number of correctly predicted detection instances versus the labelled ground truth images. Since there is only a single-class object in this study, a confusion matrix can be represented in a 2×2 matrix. (Fig 2). Out of 1473 labelled object, the model predicted 1317 of them as true positives, 137 false positives where it predicted the background as stomata and 19 false negatives where it predicted the stomata as background. This metric was taken from the confusion matrix of the training dataset.

**Fig 1.**
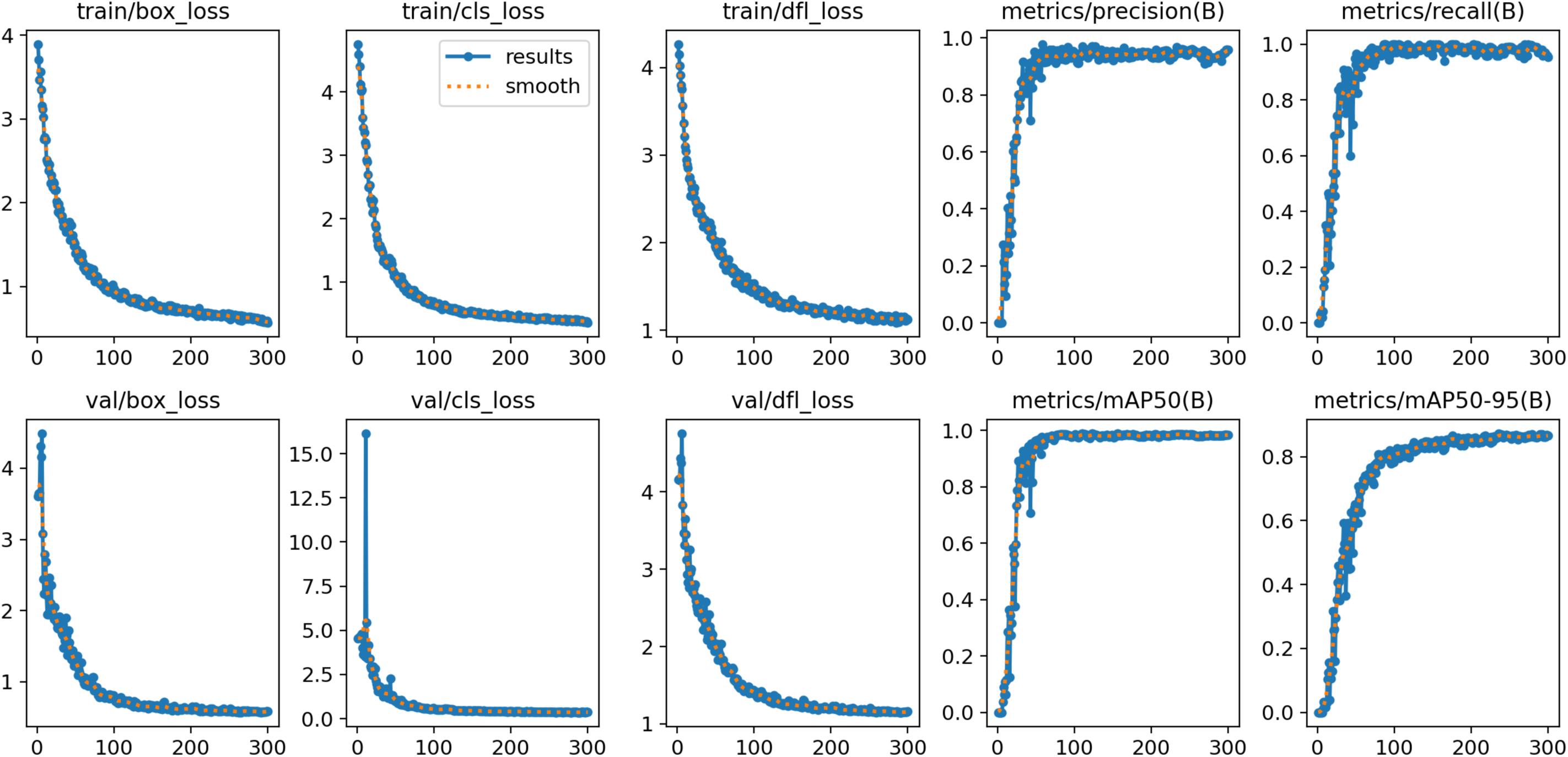
Evolutionary curves of the YOLOv12 model’s training metrics. The top panel displays the progression of box loss, class loss, and df loss, while the bottom panel shows validation performance. Accuracy metrics such as precision, recall, mAP@0.50, and mAP@0.50–0.95 approach a value of 1, indicating strong overall performance. The mAP@0.50–0.95 peaks at 0.77, suggesting the model effectively draws bounding boxes, though its precision diminishes slightly at finer resolutions near the stoma.

**Fig 2.**
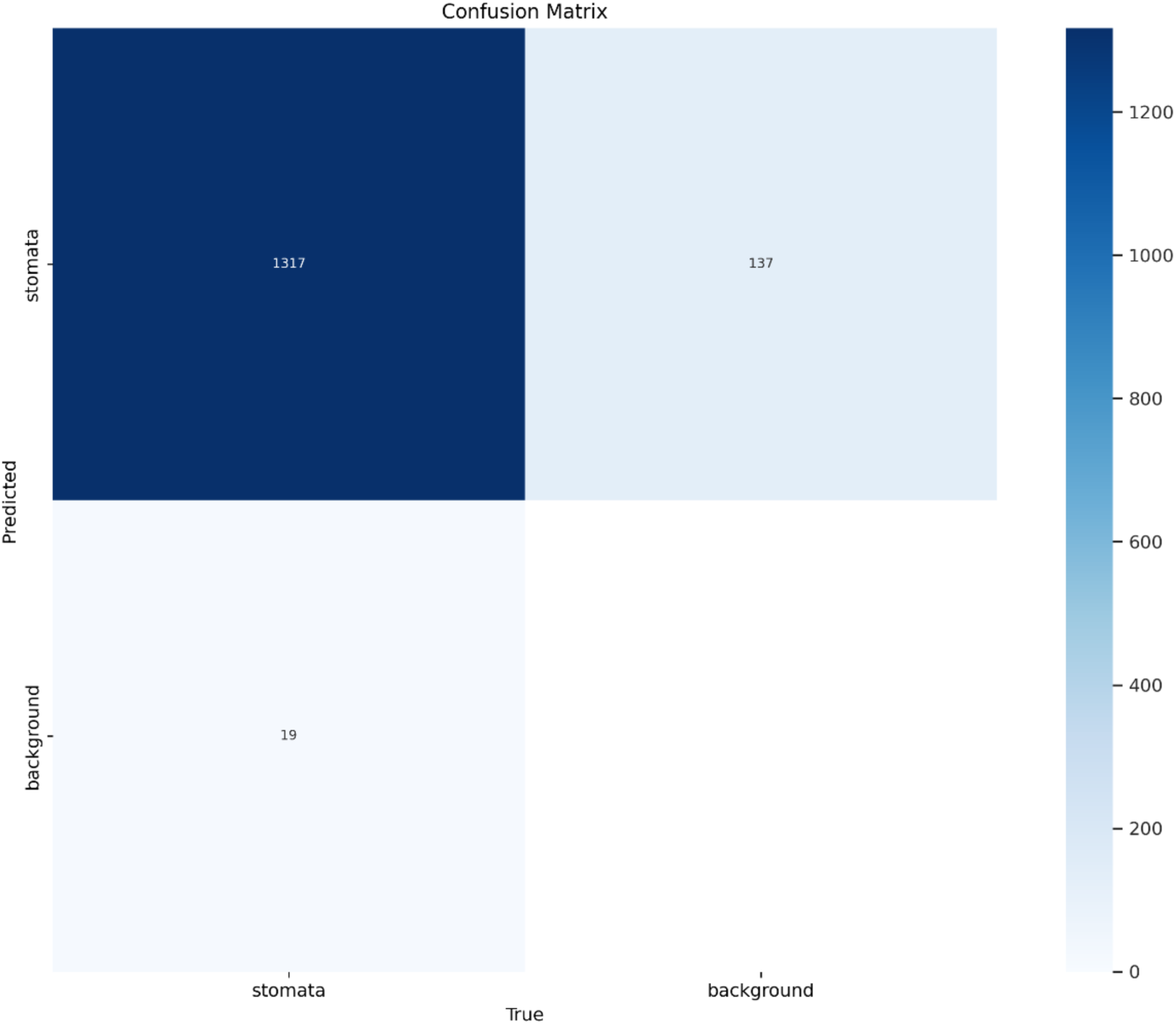
Confusion matrix comparing predicted and actual stomatal counts from the YOLOv12 test results. The top-left quadrant represents true positives, where the model accurately identifies stomata. The top-right quadrant shows false positives, indicating instances where background elements were incorrectly classified as stomata. The bottom-left quadrant corresponds to false negatives, where stomata were mistakenly labelled as background.

We then evaluated the training performance of both models. RF-DETR quickly reached its highest metric scores within the initial five epochs, whereas YOLOv12 took a minimum of 50 epochs to attain its peak performance before stabilising (Fig 3). These results suggest that RF-DETR requires lesser training cycles and training may require less computation time and resource to achieve its peak performance. Though slower in training, the final YOLOv12 model surpassed RF-DETR in all key performance metrics (Fig 4). YOLOv12 achieved a precision of 0.948 and a recall of 0.959, outperforming RF-DETR, which recorded a precision of 0.929 and a recall of 0.900. The lower recall score of RF-DETR suggests a greater likelihood of detecting false negatives where the model fails to detect some stomata even when they may be present in the image. The performance of YOLOv12 and RF-DETR models were assessed through the stomata detection task in rice, barley and sugarcane leaves. Fig 5 shows the original (left), YOLOv12 (middle) and RF-DETR (right) with detection output images displayed side-by-side, demonstrating the models’ accuracy in detecting stomata in rice, barley and sugarcane respectively. The trained YOLOv12 model can detect all stomata accurately while the RF-DETR model misses some of the stomata in our test dataset.

**Fig 3.**
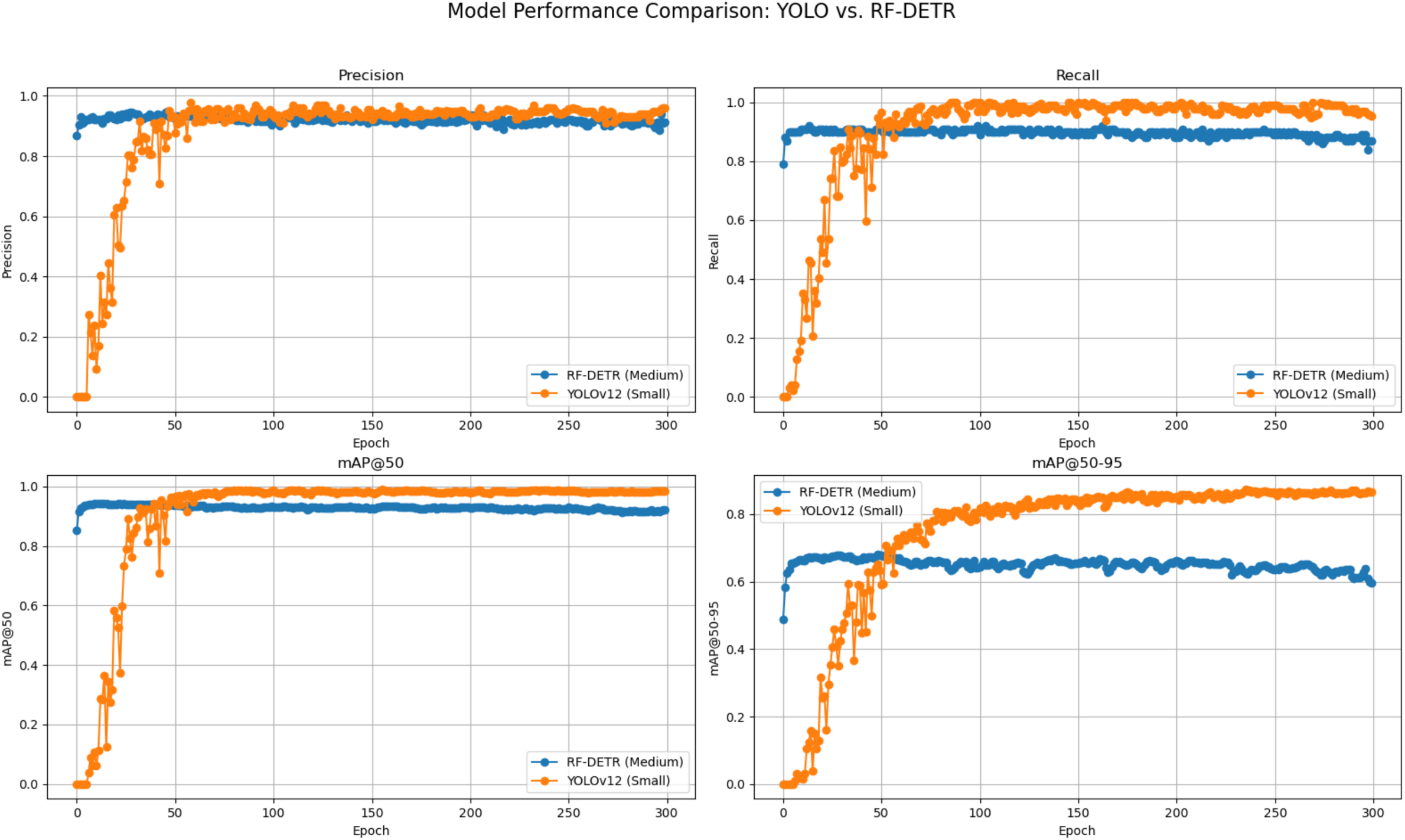
The training progression of the YOLOv12 and RF-DETR models and their accuracy metrics. YOLOv12 and RF-DETR have strong overall performance with precision scores of 0.985 and 0.939 respectively. Recall is also lower in RF-DETR with a score of 0.90 as compared to YOLOv12 with a score of 0.959. The mAP@50–95 peaks at 0.872 in YOLOv12 and 0.685 in RF-DETR. Results are summarized as bar charts in Fig 4.

**Fig 4.**
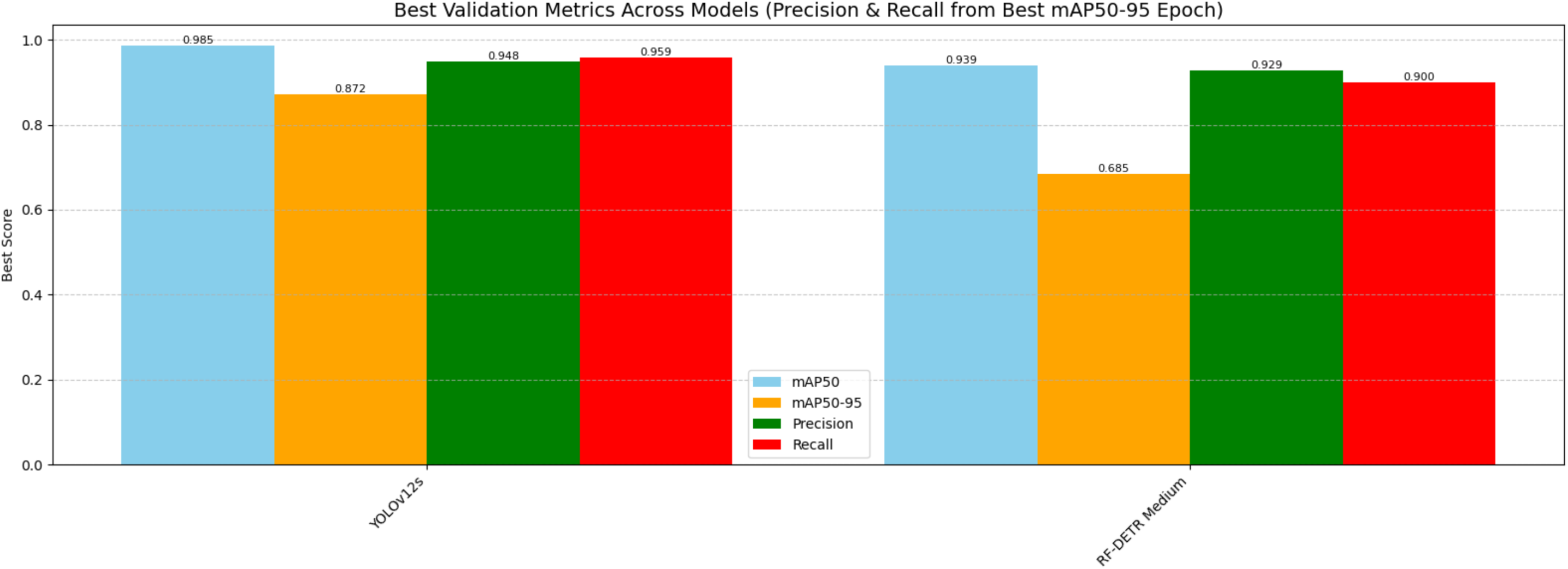
Performance metrics of mAP50, mAP50-95, precision and recall of the validation image dataset. mAP50 and mAP50-96 scores are higher in YOLOv12 at 0.985 and 0.872 as compared to RF-DETR at 0.939 and 0.685 respectively. Similarly, YOLOv12 also has a higher precision of 0.948 and recall of 0.959 while RF-DETR has a precision of 0.929 and recall of 0.900.

**Fig 5.**
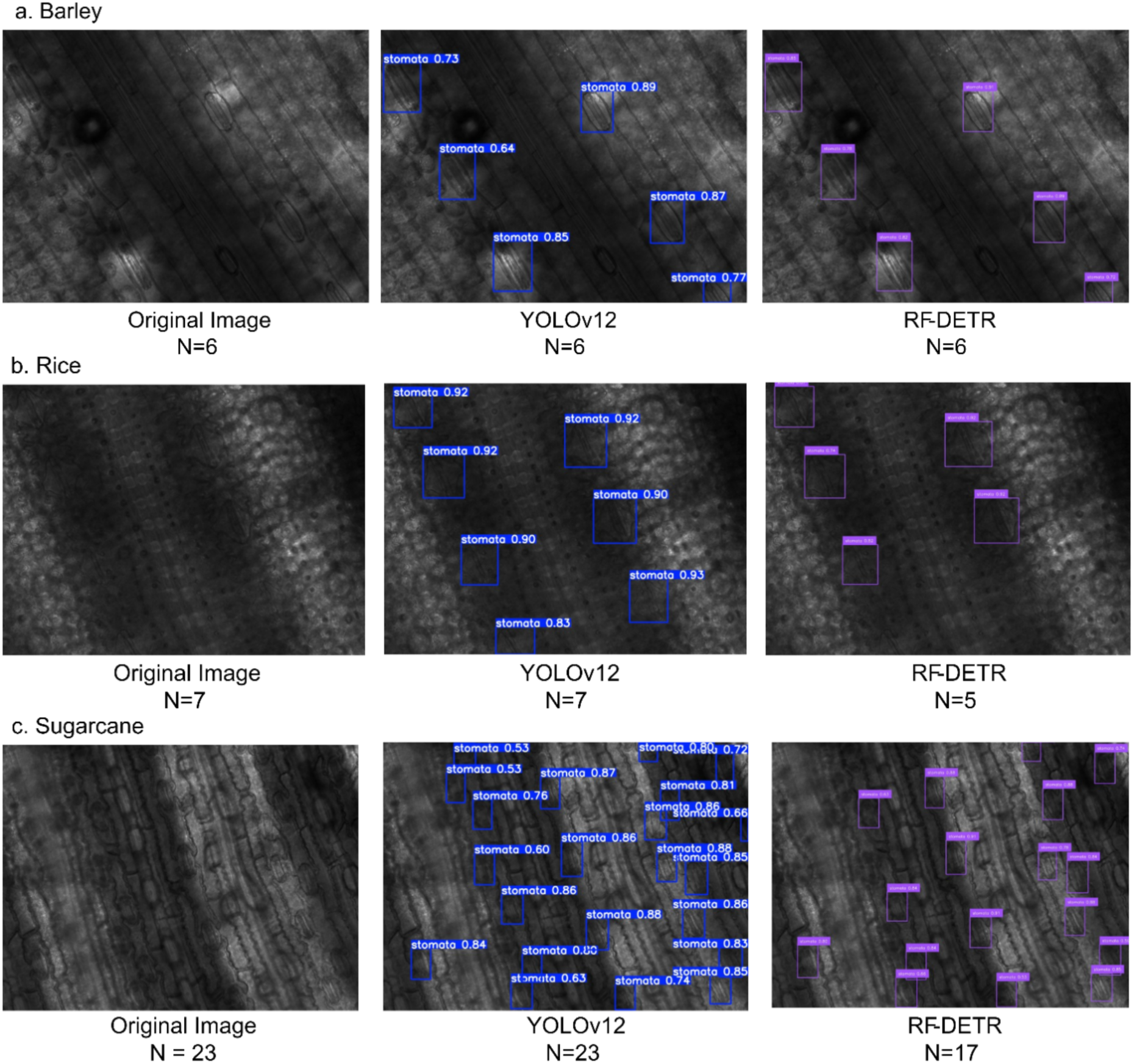
Comparative stomata detection of inference results of YOLOv12 and RF-DETR models. a. barley b. rice and c. sugarcane. YOLOv12 successfully detected every stoma present in the original images, even those that were partially obscured or only partially captured. RF-DETR underperformed and only detected stomata that were clearly imaged. N refers to the number of stomata counted.

Testing of both models was then subsequently conducted using previously unseen barley and Arabidopsis leaf images sourced from the publicly available StomaAI GitHub repository. Unlike the training dataset, the images in the GitHub repository obtained were all non-destructive leaf samples. YOLOv12 showed higher accuracy in detecting stomata, consistently producing confidence scores of 0.7 or above (Fig 6 and 7). This performance indicates that training the model on images with dark backgrounds notably improved its ability to extract relevant features, allowing it to effectively identify stomata in both light and dark grey-scale backgrounds. In comparison, RF-DETR produces bounding boxes with relatively lower confidence scores, typically in the range of 0.5 and 0.7. The model also fails to detect several stomata in some of the images (Fig 8). The model performed worse in *Arabidopsis* images as it misidentifies stoma present in background regions that have stroke-like patterns that resemble elliptical objects but with no true stomata (Fig 9).

**Fig 6.**
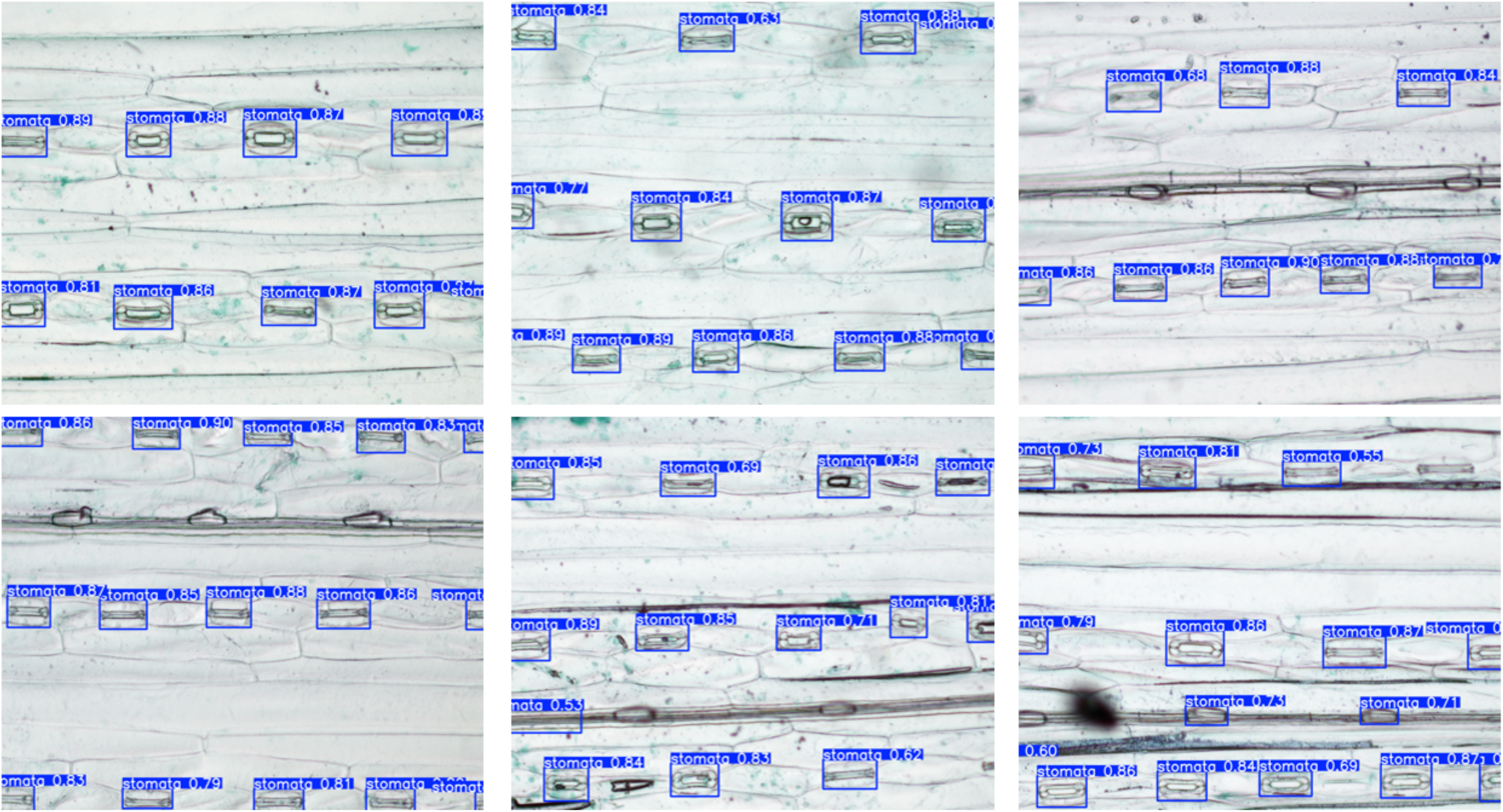
YOLOv12 inference on unseen images of non-destructive leaf images of sample barley images sourced from StomaAI GitHub repository. All stomata were detected except one stoma in the bottom right image and another vein-like structure that was misidentified as a stoma in the bottom middle image.

**Fig 7.**
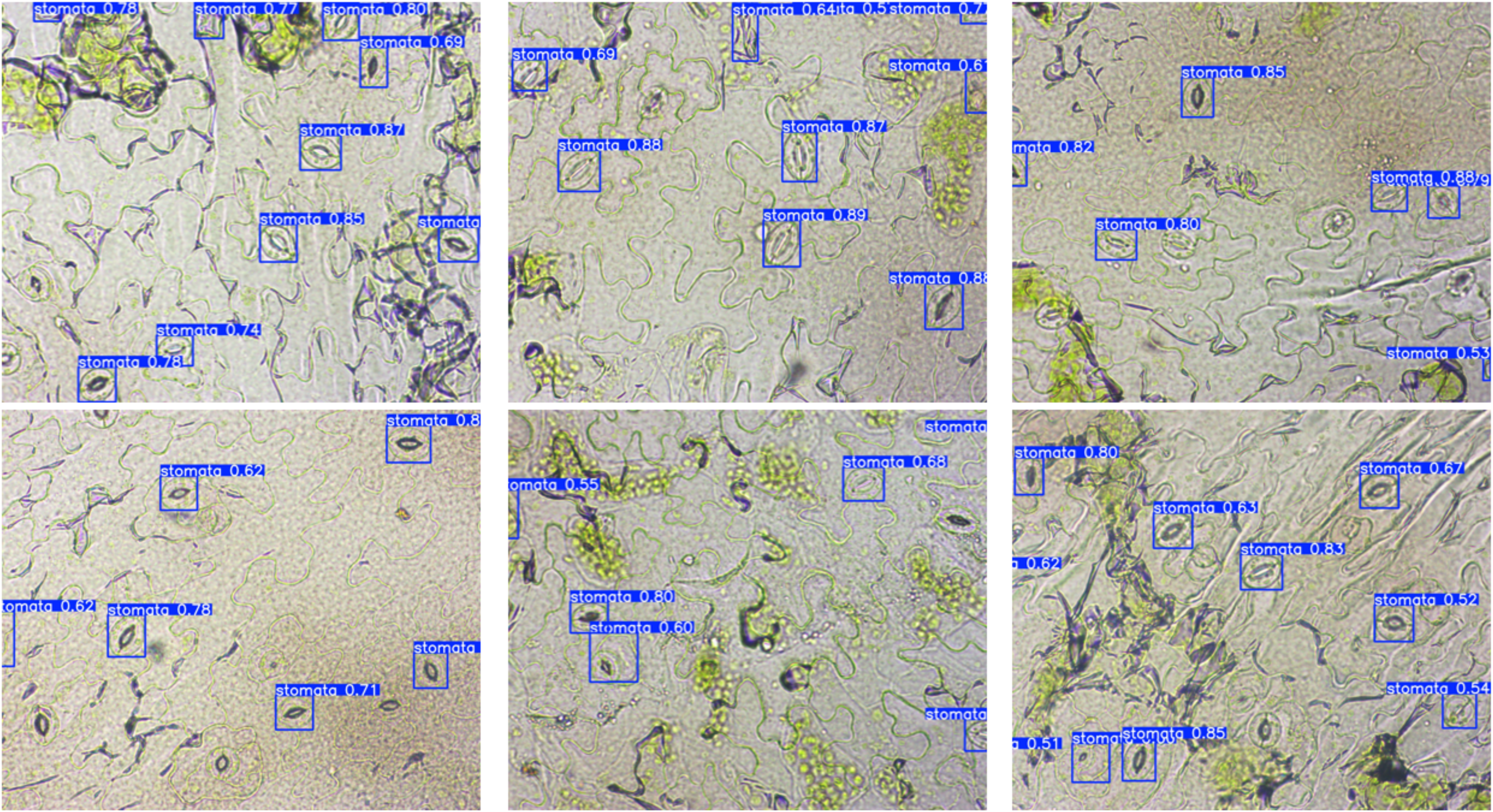
YOLOv12 inference on unseen images of non-destructive leaf images of sample *Arabidopsis* images sourced from StomaAI GitHub repository. Most *Arabidopsis* stomata were detected by YOLOv12 except two stomata in the top right image and one in the lower right image.

**Fig 8.**
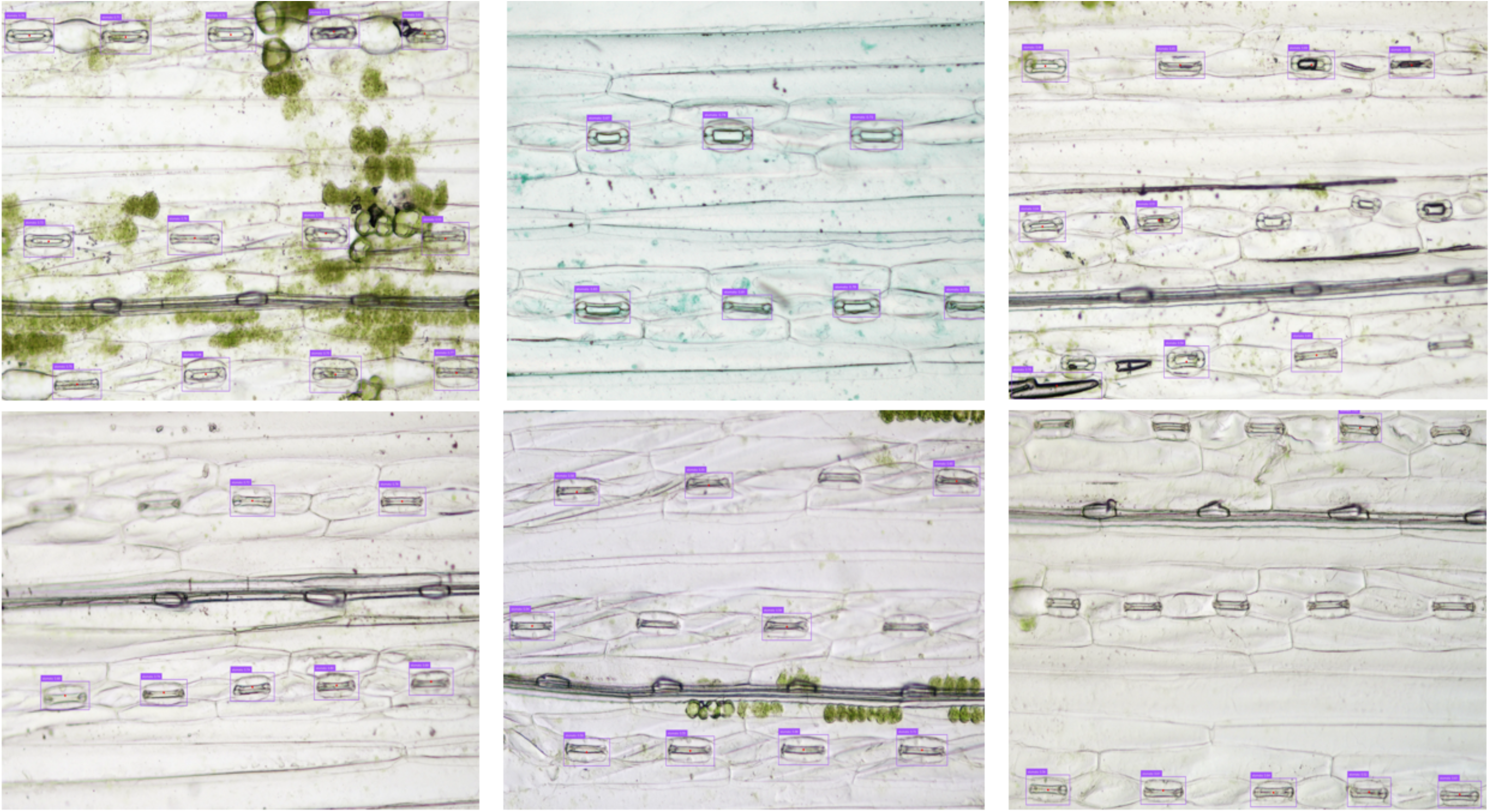
RF-DETR inference on unseen images of non-destructive leaf images of sample barley images sourced from StomaAI GitHub repository. Most stomata were detected but the model failed to detect several stomata especially in the lower right image.

**Fig 9.**
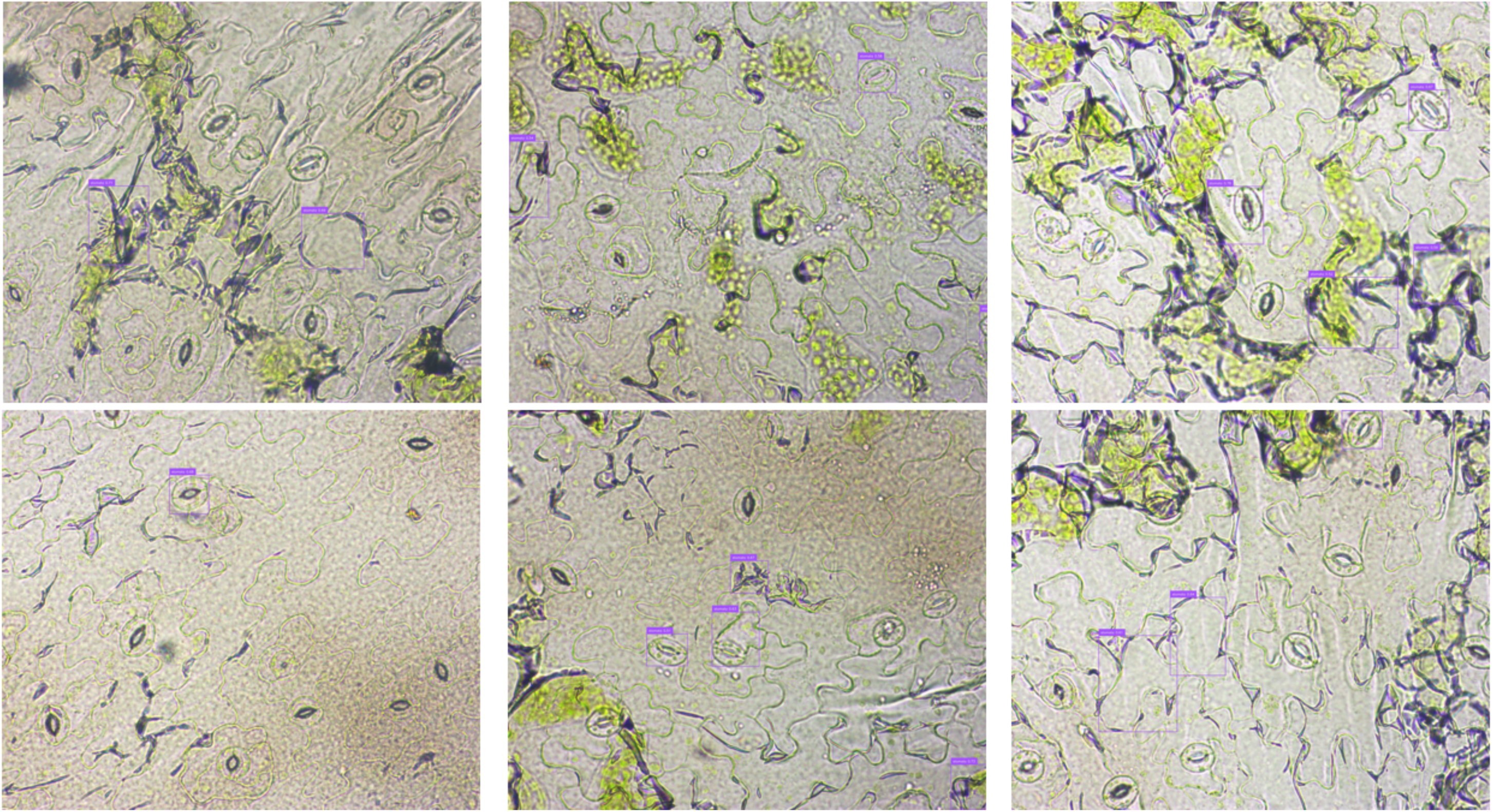
RF-DETR inference on unseen images of non-destructive leaf images of sample *Arabidopsis* images sourced from StomaAI GitHub repository. The model did not identify *Arabidopsis* stomata accurately where it drew bounding boxes around empty spaces that have some resemblance of elliptical objects in the top left and bottom right images.

In a proof-of-concept experiment, three sugarcane varieties (Var1, Var2 and Var3) were subjected to StomaQuant testing on the YOLOv12 model. Inference results in the form of annotated images from this test can be found in the supplementary data. The calibrated scale bar on the BX53 microscope indicates that the 20× magnification scale corresponds to 0.32 micrometre per pixel and 40× magnification corresponds to 0.16 micrometre per pixel. Using this calibration scale information, the stomatal traits could be calculated based on the number of pixels the segmented stoma occupies in the image and the number of predicted stomata bounding boxes.

**Area per pixel squared microns** = (microns per pixel)^2^

**Average stoma size squared microns** = (stomata pixels count × area per pixel squared microns) ÷ number of stomata counted)

**Total image area squared microns** = total image pixels × area per pixel squared microns

**Stomatal Density** = (stomata pixels count ÷ total image pixels) × 100

A total of 300 images were captured from the three sugarcane varieties and stomata traits information were inferred from the YOLOv12 model. Figure 10 shows that Var2 has the lowest stomata density, followed by Var1 and Var3. The adaxial surface tend to have lesser stomata than the abaxial side. Fig 11 shows the estimated stomata size in squared micrometre. Like stomatal density, Var2 also has smaller stomata size as compared to Var1 and Var3. No significant differences were detected between adaxial and abaxial leaf surface. Using one-way ANOVA and post-hoc analysis, we were able to deduce and conclude that Var2 has the highest drought-tolerance ability, followed by Var1 and Var3. Var2 has smaller stomata and statistically significant lesser number of stomata per squared area.

**Fig 10.**
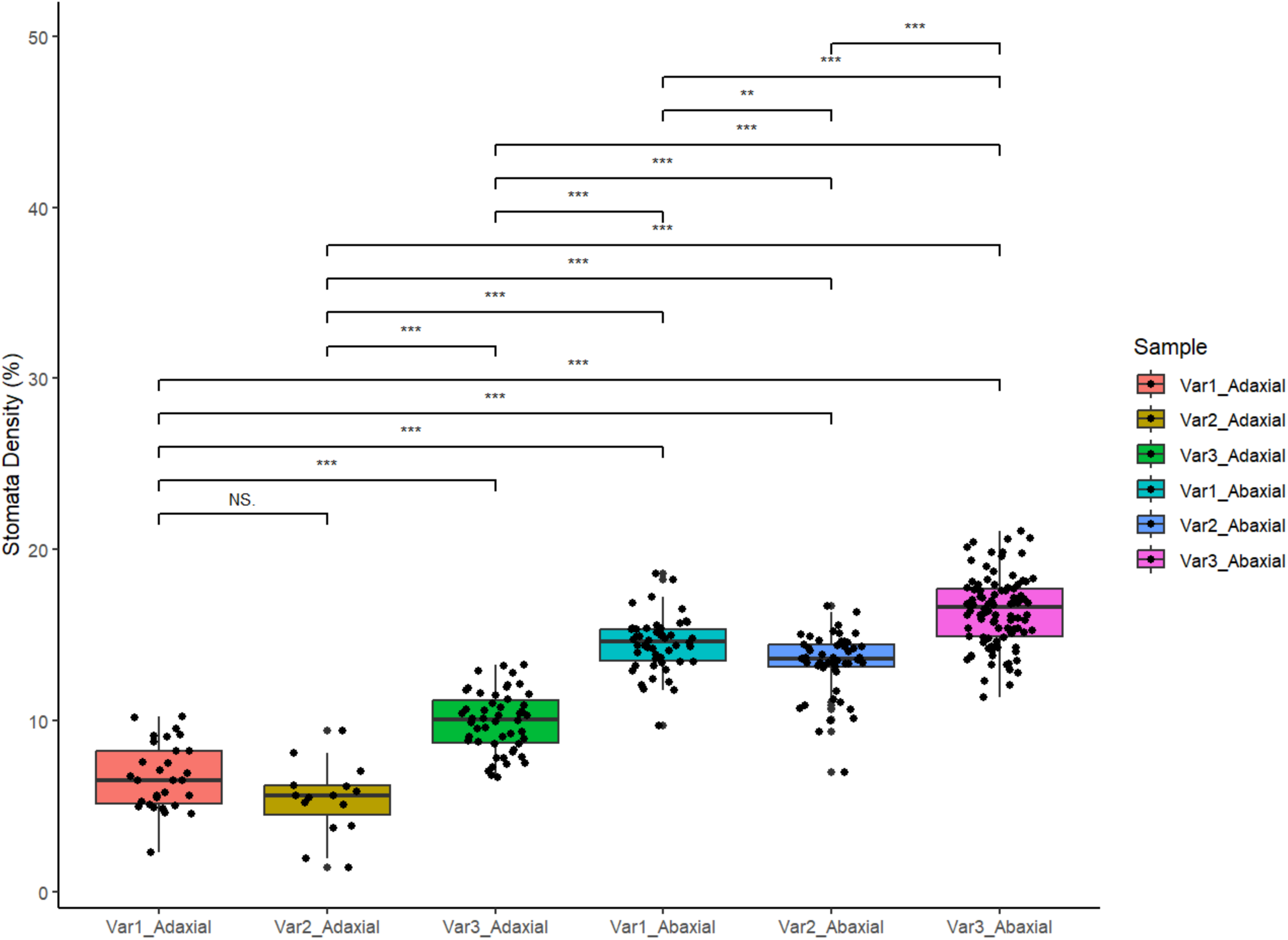
Comparative stomata density inference results of YOLOv12 on sugarcane varieties. The adaxial surfaces have lesser stomata than the abaxial surface. Var2 show the lowest stomatal density, followed by Var1 and the one with the highest stomata density in Var3. Post-hoc analysis showed that var1 and var2 have significantly lesser number of stomata than var3.

**Fig 11.**
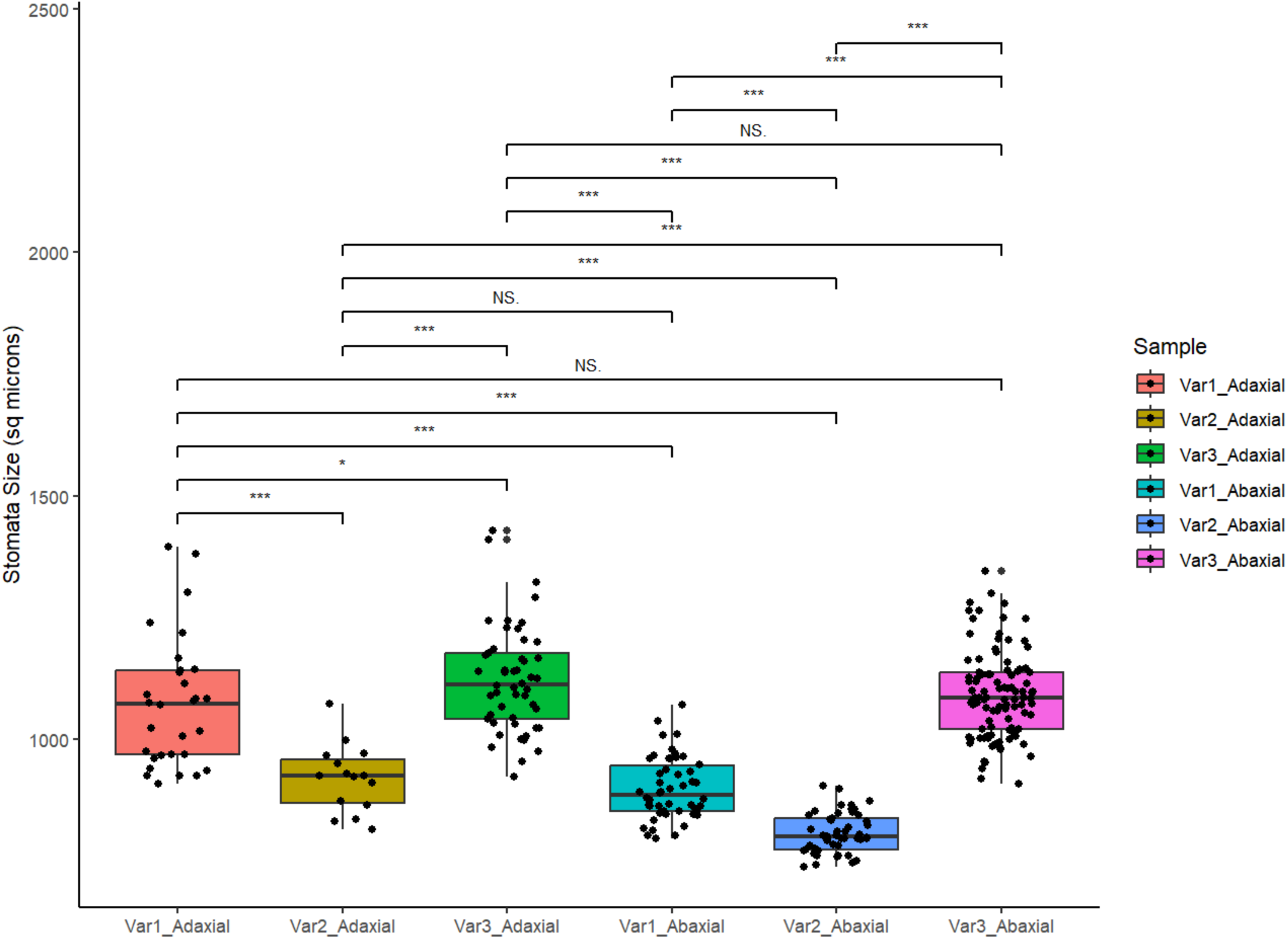
Comparative stomatal size inference results of YOLOv12 on sugarcane varieties. Stomatal size estimates showed that Var2 has the smallest stoma size of less than 1000 micrometer square (*μ*m^2^), followed by Var1 and Var3. Stomatal size did not vary significantly between adaxial and abaxial surfaces.

## DISCUSSION

The evolution of stomata detection technologies is in tandem with recent developments in computer vision, where each successive computer vision model offers increasingly faster and more accurate capabilities for real-time object detection. For instance, previous studies have investigated size estimation techniques using YOLOv8 segmentation with geometric shape fitting [22]. In this study, we used YOLOv12 to segment and estimate the stomata size and density on the leaves.

Our YOLOv12 model showed remarkable adaptability in stomata recognition across three different plants, where for instance the rhombus-shaped stoma of rice can be quite distinct from the elliptical-shaped stoma in barley and sugarcane. However, several images in our dataset contained occluded or out-of-focus stomata, resulting in blurring that may contribute to underestimations of stomatal density. Such limitations could introduce measurement bias and compromise the precision of quantitative assessments. To mitigate this issue, repeated sampling and image acquisition were performed to increase statistical power. Subsequent analysis using ANOVA and post-hoc comparisons can help discern treatment effects while accounting for sampling variability. For example, a reduced stomatal count observed on the adaxial and abaxial surface of sugarcane leaves in our sugarcane proof-of-concept experiment may serve as a phenotypic marker for drought tolerance. Comparative analysis of the sugarcane varieties revealed that Var2 has the lowest stomatal density and smallest stomatal size of less than 1000 squared microns on both adaxial and abaxial surfaces. In contrast, Var1 has moderately sized stomata and moderate stomatal density levels while Var3 has the largest stomata size and highest stomatal density. Stomata sizes did not vary significantly between adaxial and abaxial surfaces within the same variety. The stomata analyser tool suggests that among the three varieties, Var2 has most likely the highest drought tolerance ability.

Characterization of stomatal traits thus provides a valuable framework for screening cultivars with enhanced water-use efficiency (WUE). WUE refers to the ratio of carbon dioxide assimilated through photosynthesis to the quantity of water vapor released into the atmosphere [23]. Under drought conditions, levels of abscisic acid tend to rise, while hydraulic conductivity declines. This leads to a reduction in guard cell turgor pressure, causing the stomatal aperture to close, reducing stomatal conductance [24]. Several studies have performed targeted genetic modifications to understand the effect and relationship between stomata size and stomatal densities with drought tolerance in crop species. For instance, overexpression of *SDD1* gene in maize and tomato have resulted in lower stomatal density that resulted in improved WUE [25, 26]. Similarly, overexpressing *HvEPF1* in barley has also resulted in 50% reduction in stomatal density and give rise to leaves with smaller guard cells that helped the transgenic barley withstand drought stress [27]. Likewise, overexpression of *OsEPF1* gene in rice also helped improved WUE, where two transgenic rice lines showed 58-88% reduction in stomatal densities during the vegetative growth stage [28].

To date, CNNs-based algorithms continue to outperform traditional machine learning models in automated feature extraction from 2D images, demonstrating efficacy in both classification and regression tasks [29]. YOLO series model has been a popular choice due to its lightweight architecture and accurate real-time detection. In this study, we showed that the YOLOv12 model outperformed RF-DETR when tested against unseen images of barley and *Arabidopsis* sourced from https://github.com/XDynames/SAI-app. It is likely that the observed improvement in performance is attributed to the newly added attention mechanism in the YOLOv12 algorithm, adapted from transformer-based architectures. Unlike earlier iterations in the YOLO series, YOLOv12 integrates transformer-style self-attention and a Residual Efficient Layer Aggregation Network (R-ELAN) architecture [30]. This allows YOLOv12 to capture global context similarly to DETR models while maintaining the strengths of CNNs for local feature extraction of small repetitive structures like stomata. In comparison, transformer-based DETR models generate predictions using bipartite matching via an encoder-decoder architecture, where it learns about the relations of objects and the global image context to create a set of predictions in parallel [31]. RF-DETR has a DINOv2 pre-trained model as its backbone and it converts the image and stomata annotations into a string of sequence tokens utilising transformer to capture long-range dependencies between input tokens [32]. Implementing transformer self-attention has been a challenging problem in computer vision in the past due to slower speeds as compared to CNN-based model as the computational space needed to process image patches as tokens is significantly larger than a string of words in multi-head self-attention calculations [33]. To reduce the computational space complexity, newer models like RF-DETR incorporate parallel computing strategies where attention can be computed in a hierarchical manner at different scales to divide the feature map into smaller manageable grids with residual connections to fuse the hierarchical outputs. This allows DETR models to adjust the attention computation dynamically and speeding up convergence during training [34]. It is also evident in our training process as seen previously in Fig 3 showing that RF-DETR converges within the initial five epochs while YOLOv12 converges only after 50 epochs. Additionally, Fig S2 also shows that RF-DETR model converges early, and performance of model also decreased beyond epoch 50 likely due to overfitting. Results indicate that the model tends to memorize the training data, which may lead to suboptimal generalization on unseen datasets. The latest YOLOv12 bears resemblance to the DETR models and is built with area attention and 7×7 convolutional layers architecture coupled with R-ELAN links that harness the strengths of transformer-based global relations learning and preserve the core YOLO-series CNN-based feature extractions for real-time object detection. Our results indicate that YOLOv12 demonstrates greater sensitivity and accuracy in stomata detection, although it requires more training epochs to reach its peak performance.

## CONCLUSIONS

In summary, YOLOv12 and RF-DETR models can be used for stomata detection task. Based on our dataset and training parameters, the YOLOv12 model perform better at recognising stomata from images of rice, barley, sugarcane and *Arabidopsis* leaves derived from both destructive and non-destructive methods. High-throughput image processing and post-hoc statistical analyses can be implemented with larger sample sizes in plant breeding programs. Our stomata analyser tool can assist plant breeders to screen and select for drought-tolerant plant varieties with smaller stomata size and lower stomatal densities.

## DATA AVAILABILITY

The training, validation and test images are publicly accessible for download via Dryad repository at 10.5061/dryad.3j9kd51zr.

(http://datadryad.org/share/LINK_NOT_FOR_PUBLICATION/A7j9OfWGyGwgBGwxWMImNfn_XLovoCEFWG2ccCEARQQ)

All python scripts used in this work are available in the GitHub repository at https://github.com/kjxlau/StomaQuant.

The web application is hosted at https://huggingface.co/spaces/kennylau91/stoma.

## ACKNOWLEDGMENTS

We thank the Temasek Life Sciences Laboratory, Singapore for research funding and computational support. We thank the Fungal Patho-biology group for helpful discussions and suggestions; and GMP Indonesia for sharing the sugarcane varieties.

## SUPPLEMENTARY INFORMATION

**Fig S1.**
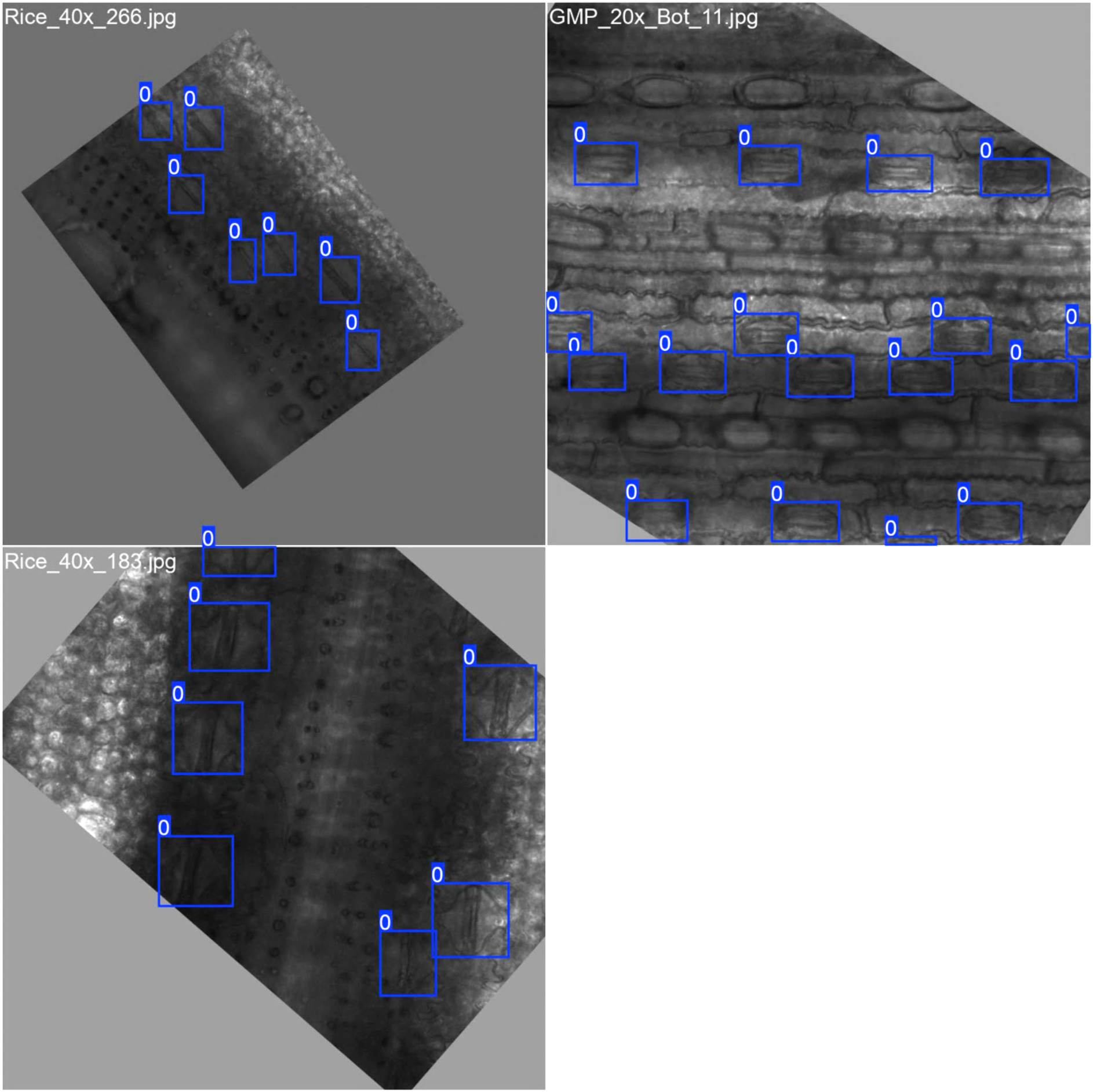
Image augmentation images used for model training. A scale of 0.5, vertical flip of 0.5, horizontal flip of 0.5, and rotation of 90 were applied to generate more images for model training. The figure displays three images with bounding boxes generated during image augmentation prior to training.

**Fig S2.**
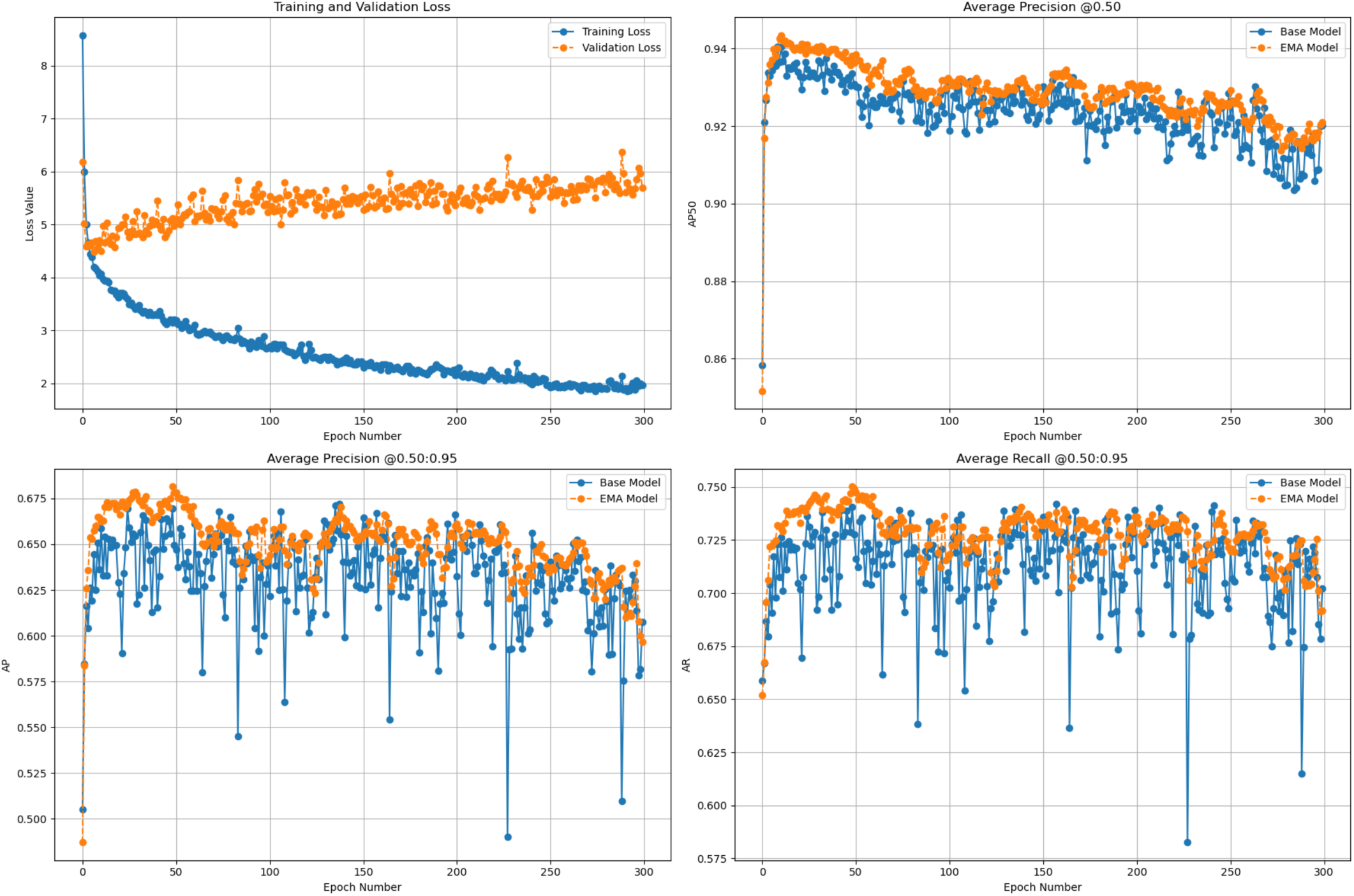
Training metrics and performance of RF-DETR model. Training loss curve shows that the model is learning by minimizing the error on the training set. However, the increase divergence between training and validation loss suggests overfitting when model is trained to epoch 300. Performance of RF-DETR model peaks around epoch 20 at 0.94. Exponential moving average (EMA) beats the base model for both precision and recall with overlapping threshold of 0.50-0.95, yielding scores of 0.68 and 0.75 at epoch 50.

